# Investigating the unknown functions in the minimal bacterial genome reveals many transporter proteins

**DOI:** 10.1101/381657

**Authors:** Magdalena Antczak, Martin Michaelis, Mark N Wass

## Abstract

The recent identification of the minimal bacterial genome revealed that nearly one third (149) of the 473 encoded genes were of unknown function, demonstrating our limited understanding of the essential functions of life. Application of state of the art *in silico* methods for functional annotation demonstrated that these proteins of unknown function lack orthologs, known protein domains, and templates to model their structure. Combination of the results from different complementary approaches enabled functions to be assigned to 94 of the 149 proteins, although often with general terms such as transporter or DNA binding. 22 likely transporter proteins were identified indicating the importance of nutrient uptake into and waste disposal out of the minimal bacterial cell, where many metabolic enzymes have been removed. These results advance our understanding of the minimal bacterial genome and therefore aid synthetic biology and its application to biotechnology.

## Introduction

A long-term goal of synthetic biology has been the identification of the minimal genome, i.e. the smallest set of genes required to support a living organism. This has been a systematic process using species of Mycobacteria, which have some of the smallest bacterial genomes. Initial sequencing of *Mycoplasma genitalium* identified a genome containing 525 genes and comparison with the *Haemophilus influenza* genome (1815 genes) suggested a common core of 256 genes as a possible minimal set of genes essential for life.^1^ Subsequent global transposon mutagenesis experiments suggested this minimal set was larger containing 375 genes.^2^

The bacterium with the smallest genome generated to date is based on the faster growing *Mycoplasma mycoides*. Its minimal bacterial genome consists of 473 genes including essential genes and a set of genes associated with growth, termed ‘quasi-essential’.^3^ The minimal genome study assigned function to genes from the minimal genome by considering matches to existing protein families in the TIGRfam^4^ database, genome context and structural modelling.^3^ Proteins were annotated with molecular functions and grouped into 30 biological process categories (including an unclear category, where the biological process was not known). The proteins were further grouped into five classes according to the confidence of the functional annotations that they had been assigned. The five classes were: *equivalog* (confident hits to TIGRfam families), *probable* (low confidence match to TIGRfam families supported by genome context or threading), *putative* (multiple sources of evidence but lower confidence), *generic* (general functional information identifiable e.g. DNA binding or membrane protein, but specific function unknown) and *unknown* (unable to infer even a general function). The final two confidence classes, *unknown (65 genes)* and *generic* (84 genes) form a group of genes whose function is unknown. Hence, almost a third (149) of the encoded 473 proteins are of unknown function, which emphasises our limited understanding of biological systems.^3^

This lack of functional annotation is not restricted to the minimal bacterial genome. Recent experimental approaches have begun to identify the function of ‘hypothetical’ proteins of unknown function.^5^ However, the continual improvement of high throughput sequencing methods has resulted in a rapid increase in the number of organisms for which genome sequences are available and the functional annotation of the encoded gene products lags behind.^5^ Less than 1% of the 120 million protein sequences in UniProt^6^ are annotated with experimentally confirmed functions in the Gene Ontology (GO)^7^ (as of July 2018). To address this gap, computational methods for protein function prediction have been developed and significantly advanced over the past 15 years as demonstrated by the recent Critical Assessment of Functional Annotation (CAFA) challenges.^8,9^

Here, we performed an extensive *in silico* analysis of the proteins of unknown function encoded by the minimal bacterial genome using an approach that combined sixteen different computational methods ranging from identification of basic properties (e.g. protein domains, disorder and transmembrane helices) to state of the art protein structural modelling and methods that infer GO based protein functions, including those that have performed well in CAFA experiments. This enabled us to assign functions to 94 of the 149 the previously uncharacterised proteins.

## Results

Initially, the basic properties of the proteins encoded within the minimal bacterial genome were characterised. This considered the presence of orthologs, domain architecture, protein disorder, transmembrane domains, and the ability to model protein structure. The properties of the proteins in the five different confidence classes *(equivalog* to *unknown)* were compared to identify if there were differences between the proteins in each of the confidence groups that may relate to the ability to infer their function.

### Orthologs for the proteins in the minimal genome

Hutchison et al.^3^ used BLAST to identify homologs of the minimal genome proteins in a set of 14 species ranging from non-mycoides mycoplasmas to Archaea. They found that while many of the proteins from the *equivalog, probable, putative* and *generic* classes have homologs in all 14 species, very few of the sequences in the unknown class had homologs in the 14 species outside of *M. mycoides*, with none in *M. tuberculosis, A. thaliana, S. cerevisiae* and *M. jannaschii*.

Here, eggNOG-mapper^7^ (See methods) was used to identify orthologs for the minimal genome proteins across the three kingdoms of life. Overall the analysis showed that very few of the *unknown* class of proteins (7%) have related sequences in eukaryotes or archaea (6%) while just over half (55%) have orthologs in other bacterial species, primarily in terrabacteria, the clade that *M. mycoides* belongs to (Figure 1A, Figure S1, Table S1). In contrast, many of the proteins in the other confidence classes have orthologs across the three kingdoms (Figure 1A, Figure S1). For example, 63%, 59% and 95% of the proteins in the *generic* class have orthologs in Eukaryotes, Archaea and Bacteria respectively (Figure 1A, Figure S1), rising to 91%, 70% and 99% for the *equivalog class*. Only two proteins from the *unknown* class had many orthologs in both Eukaryotes and Archaea. These proteins MMSYN1_0298 and MMSYN1_0302 were classified by Hutchison et al. into the Unclear and Cofactor transport and salvage functional categories, respectively. Our analysis revealed that MMSYN1_0298 is likely to be a ribosomal protein from the family L7AE and MMSYN1_0302 an oxygen-insensitive NAD(P)H nitroreductase, rdxA, both of which are functions widespread across the kingdoms of life (Tables S1-S7).

**Figure 1.**
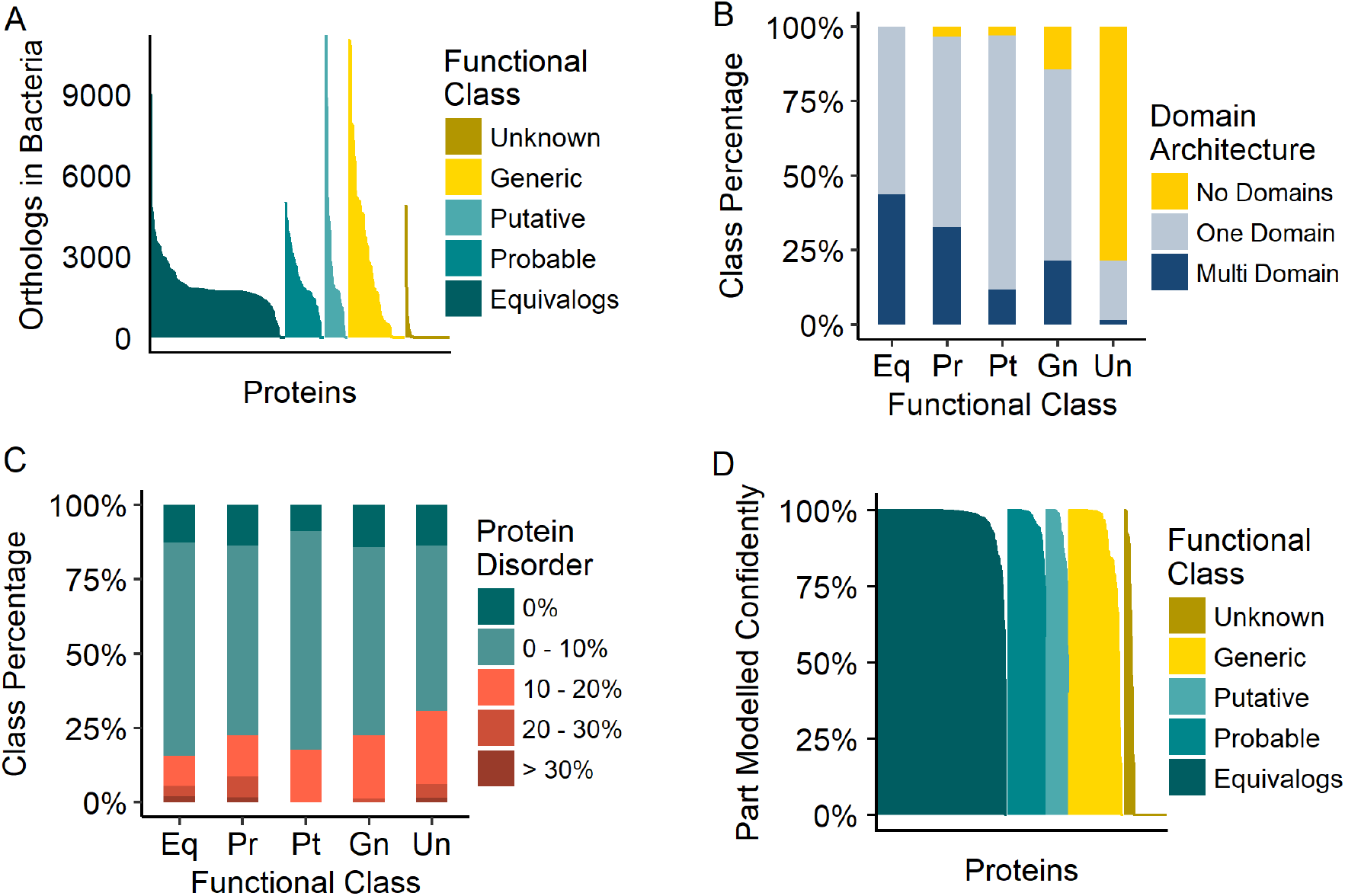
Basic characterisation of proteins in the minimal bacterial genome. A) Orthologs identified in bacteria b) proteins domains present in the minimal genome split into functional confidence *(unknown, generic, putative, probable* and *equivalog* c) predicted protein disorder in the minimal genome proteins d) percentage of protein structure that can be confidently modified by Phre2.

### Domain Architecture of Minimal Genome Proteins

Domain analysis, using Pfam^10^ (Table S2), revealed that very few (22%) of the proteins in the *unknown* class contain known domains, which is significantly less than for any of the other four classes (Figure 1B; p < 8.3e-12; Mann-Whitney-Wilcoxon test). In contrast, all proteins in the *equivalog* class contain at least one domain and nearly half of them (44%) have a multi-domain architecture (Figure 1B). Whereas multiple domains are present in 21% of the proteins in the *generic* class and only a single protein in the *unknown* class (Figure 1B). The proteins in the *unknown* class are also clearly different to those in the *generic* class, where a domain is present in 86% of the proteins. This highlights that the *unknown* class really represents proteins that are totally uncharacterised. Further, the proteins in the *unknown* class also have more disordered regions (see methods) than the other groups (Figure 1C), although this does not reach statistical significance (p > 0.05; Chi-Square test for categorical data).

### Structural Modelling of the minimal genome

Hutchison et al., used threading to support functional assignment from TIGRfam maches.^3^ Here the Phyre2^11^ protein structure prediction server was used to model the structures of the minimal genome proteins and in turn the models generated for the proteins in the *generic* and *unknown* classes were used to infer their function. With the exception of the *unknown* class, high confidence structural templates were identified for the vast majority of proteins for at least part of the sequence (Supplementary Figure 2, Table S3). The proportion of proteins in each confidence class that could be accurately modelled was considered by identifying those for which at least 75% of the protein sequence could be modelled with a structural model confidence score (from Phyre2) of at least 90%. In the *unknown* class this applied to only nine proteins, whereas nearly all proteins in the four other confidence groups could be successfully modelled (Figure 1D). Again, this demonstrates differences between the *unknown* and *generic* classes, while most of the *generic* class can be modelled, this was possible for only very few proteins in the *unknown* class (Figure 1D).

### Identifying Transmembrane Proteins

Prediction of transmembrane helices revealed that the proteins in the *unknown* and *generic* classes are enriched with transmembrane proteins with 49% and 32% of their proteins predicted to have one or more transmembrane helix respectively (Figure 2A, Table S4). This contrasts with very few transmembrane proteins identified in the *equivalog* and *probable* classes (6% and 12% respectively), while 30% of the proteins in the *putative* class are transmembrane proteins (Figure 2A). These results suggest that many of the proteins that have unassigned functions may be associated with membranes. For example, 24 proteins in the *generic* class are predicted to contain six or more transmembrane helices (Figure 2B), many of which are likely to be transporters of essential nutrients from the media (see below).

**Figure 2.**
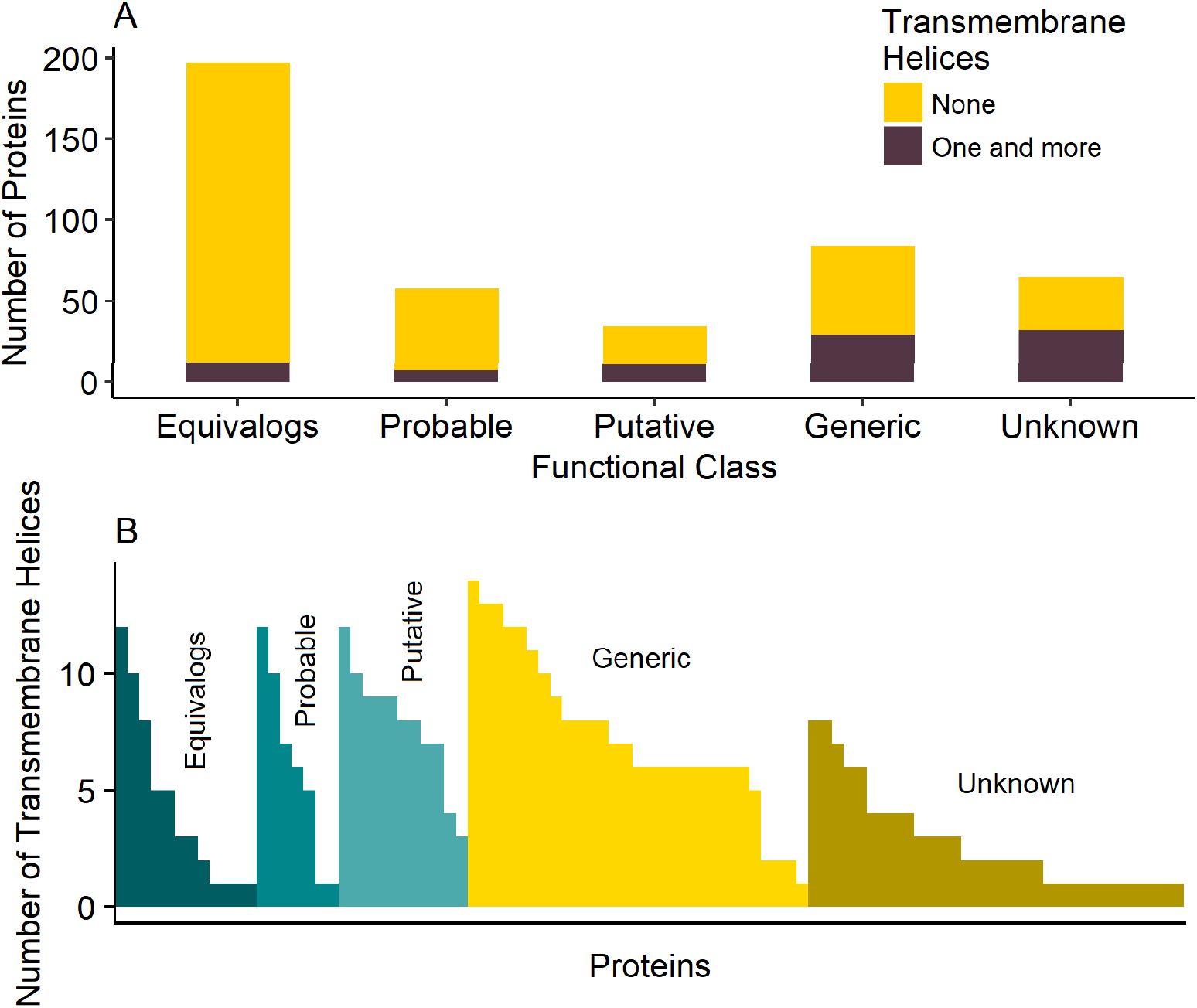
Transmembrane proteins encoded by the minimal bacterial genome. A) The number of proteins predicted by TMHMM to have transmembrane helices. B) The number of transmembrane helices present in each of the proteins in the minimal genome that is predicted to have one or more transmembrane helix.

### Inferring Protein Function for the Minimal Genome

Hutchison et al.^3^ used hits to TIGRfam, genome context and threading to functionally characterise the proteins encoded by the minimal genome. However, 149 out of 473 proteins were classified with low confidence in the *unknown* and *generic* classes. Here we sought to infer annotation for these proteins and also increase the confidence of any existing annotations.

Many different methods have been developed to predict protein function using properties ranging from protein sequence to interaction data and predicting features ranging from subcellular localisation to Gene ontology (GO) terms and protein structure.^12^ Where available as either a webserver or for download, we applied the top performing methods from the recent CAFA assessments and other established methods (see methods) to predict functions for the proteins in the *generic* and *unknown* classes. Briefly we considered, a group of methods that predicted either protein structure, domains, GO functional terms, or whether proteins are transporters (see methods). Overall functional inferences were made by manually investigating and combining the predictions and their consistency with genes from the same operon.

For nearly two thirds of the proteins (94 of 149) either a new function was identified or confidence increased in the annotation previously assigned (Figure 3, Table S5). This included 58 proteins where we identified new functional information. 29 of these proteins had previously been classed in the *unclear* functional category. The majority of these proteins were predicted to have transporter related functions, with 22 proteins added to the 84 already in this functional category (Figure 3A). Further, one protein was assigned to the cytosolic metabolism category, three to the preservation of genetic information category, and three to the expression of genetic information category (Figure 3A).

**Figure 3.**
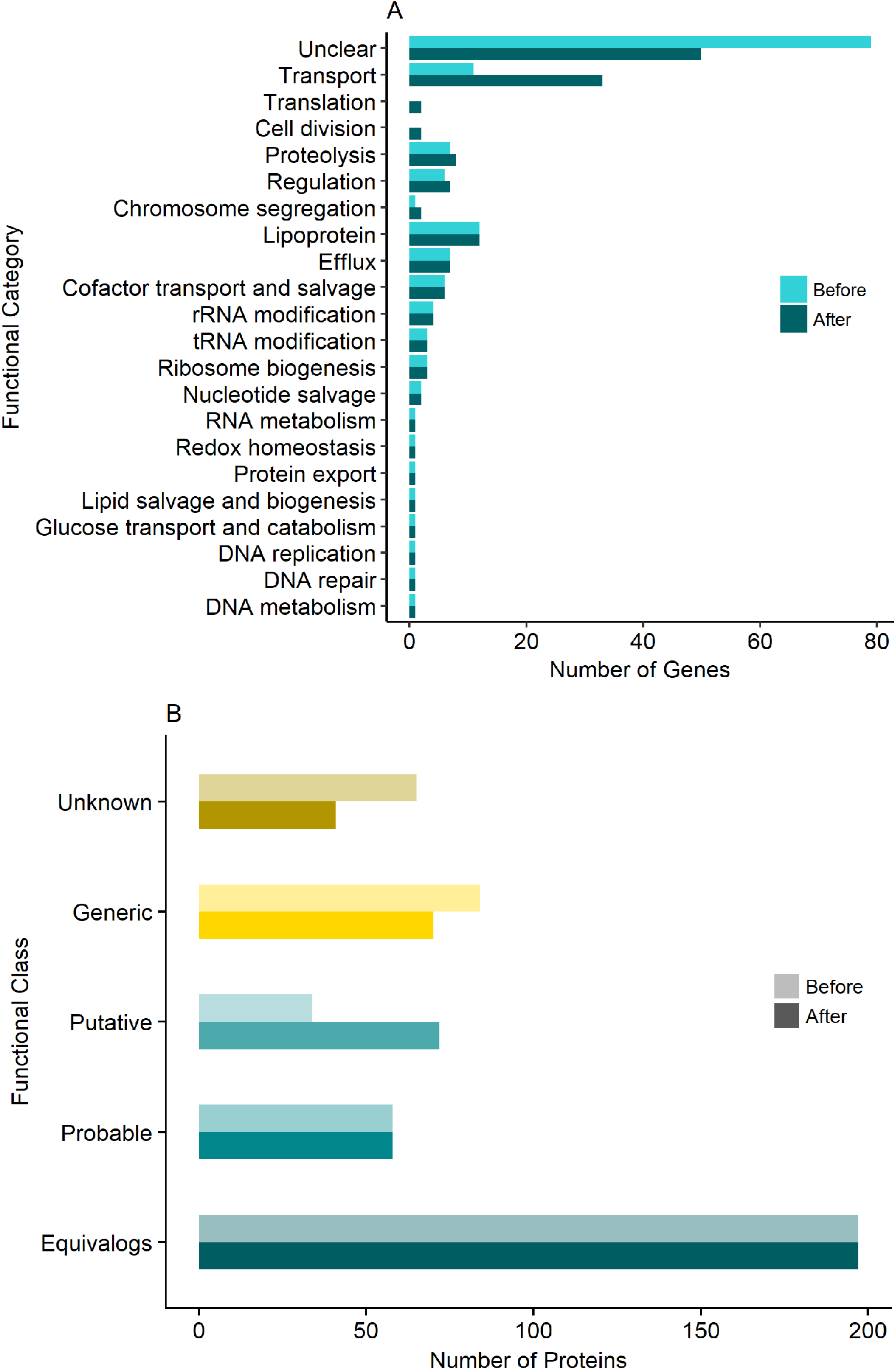
Functional annotations of the minimal bacterial genome. The number of proteins in each of the A) protein biological process categories and B) functional confidence classes is shown with the original minimal genome annotation and the annotations identified here. C) Shows the change in functional confidence classes, coloured based on original group.

For 59 proteins the confidence class was increased. 23 proteins were assigned additional functional annotations, while for the remaining 36 proteins the confidence in the existing annotation was increased. This included 24 proteins that were upgraded from the *unknown* class to either the *generic* (21) or *putative* (3) classes. For a further 35 proteins the confidence class was upgraded from *generic* to *putative*. The remaining 90 proteins (41 *unknown* and 49 *generic)* remained in the same evidence class. Despite additional functional information, there was not sufficient data to increase the confidence of the annotation for these proteins.

For some proteins while there appeared to be evidence for a given function from multiple sources but on closer inspection it was difficult to assign more confident annotations (Figure S3). For example, MMSYN_0138, is homologous to the ATP binding region of ABC transporters but the ATP binding site is not conserved, which casts some doubt on this function (Figure S3A). For MMSYN_0615, matches from four methods suggest a Phenylalanine-tRNA ligase function. This includes GO term predictions (combined from multiple methods), structural matches based on the beta chain of bacterial Phenylalanine-tRNA ligases from both Phyre2 and Gene3D/CATH and a hit to the TIGRFam Phenylalanine-tRNA ligase β subunit family (Figure S3B). However, MMSYN_0615 only contains 202 residues and the beta chain of bacterial Phenylalanine-tRNA ligases contain closer to 800 residues, making it unlikely that MMSYN_0615 performs this function and there are many more predictions that agree with the existing tRNA binding protein annotation (Figure S3B).

While functional annotations have been inferred for a considerable proportion of the proteins of unknown function, there remain 50 proteins for which the biological process they belong to is still classed as unclear. 28 of these proteins are also in the *unknown* confidence class, while 22 are in the *generic* class (four of which moved from the *unknown* to the *generic* class in our analysis) and have basic functional information such as Cof-like hydrolase, ATPase AAA family, or DNA-binding protein. Fourteen proteins have completely unknown function and remain labelled as hypothetical. These proteins do not contain any known domains or transmembrane helices, none have orthologs in other kingdoms of life and only a few within bacteria. So it seems that either these are highly species specific proteins that perform an important function within Mycobateria or they have diverged significantly such that the sequence relationships are not detected.

### Predicted Transporters and Transmembrane proteins

Transmembrane helices were identified in 41% (61) of the proteins in the *unknown* and *generic* and classes (Figure 2, Table S4). Seventeen transmembrane proteins which were not categorised as transporters were annotated with functions in cell division (1), chromosome segregation (1) and proteolysis proteins (4). However, 11 of them still remain in the unclear functional category. Our analysis suggests that 44 of the 61 predicted transmembrane proteins are likely to be responsible for membrane transport (Table S4,S5). Of the 44, 23 were previously annotated by Hutchison et al. with a range of functions including ABC transporters, S component of ECF transporters, amino acid permeases, lipoproteins, and putative magnesium-importing ATPase. Our analysis supported all of the existing transporter functions, and identified new transporters among those listed as membrane and hypothetical proteins. Of the 22 newly proposed transporters (previously listed as hypothetical or with minimal information e.g. membrane protein), five gained specific transporter functions. All five were previously classed as membrane proteins and have now been annotated as transporters; two possible ABC transporters (MMSYN_0138, MMSYN_0411), one S component of an ECF transporter (MMSYN_0877), and two belonging to the Major facilitator superfamily (MMSYN_0325, MMSYN_0881)(Tables S1-S6).

The remaining 17 proteins annotated as transporters had previously either been annotated as membrane or hypothetical proteins and we propose that these are very likely transporters but it was not possible to assign them to a specific family/type of transporter or to identify a substrate. More detailed annotations were added to some of those already identified to have a role in transport including four proteins (MMSYN1_0034, MMSYN1_0399, MMSYN1_0531, MMSYN1_0639) that were classed as FtsX-like permeases having previously been given generic transport related annotations (e.g. permease or efflux protein). For 16 others the confidence of the existing annotation was increased, for example two operons encoding proteins that transport oligopeptides (AmiABCDE – MMSYN_0165, MMSYN_0169 and PotABCD MMSYN_0195, MMSYN_0197) were moved from the *generic* to the *putative* confidence class based on support from most of the methods (Table S5).

Fifteen proteins that have been assigned transporter functions do not have transmembrane domains. They include mainly lipoproteins and ATP-binding units of ABC transporters, most of which (14) were already identified by Hutchison et al. as having membrane associated functions.

One of the two proteins proposed to be members of the Major facilitator superfamily, MMSYN_0325, was previously classified as a membrane protein (Figure 4). In agreement, the transmembrane helix prediction tool TMHMM^13^ predicted 13 transmembrane helices in the protein. Further, the structure was confidently modelled by Phyre2, with very high confidence of >98% for 26 independent structural templates, all of which had transporter functions (including members of the MSF superfamily). Gene3D^13^ (Table S7) also identified matches to an MFS general substrate transporter like domain (lactose permease functional family). Supporting this SCMMTP^14^ predicted the protein to be a transporter and function prediction methods predicted a range of transporter related functions, including symporter activity (GO:0015293), cation transmembrane transporter activity (GO:0008324) and substrate-specific transmembrane transporter activity (GO:0022891) with probabilities greater than 90% (Figure 4 and Table S6). While 3DLigandSite predicted heme and zinc binding through matches to Bacterioferritin (for the heme), it seems unlikely that these are relevant to the transporter function.

**Figure 4.**
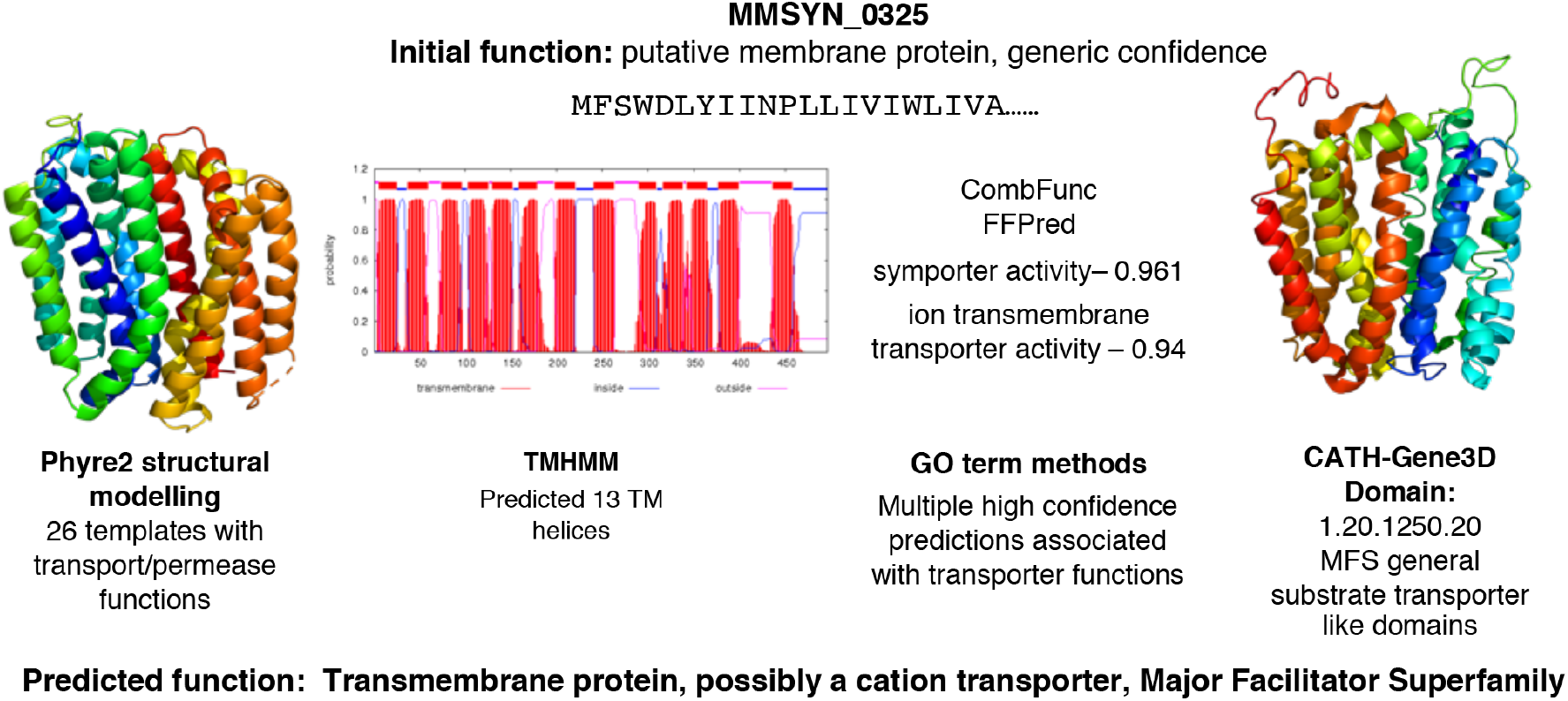
MMSYN_0615 is predicted to be a transporter and member of the Major Facilitator Superfamily. The results from Phyre2, TMHMM, the combination of GO term prediction methods (numbers shown are probability associated with each function) and CATH/Gene3D. All of these methods supported a transporter function with Phyre2 and CATH/Gene3D confidently identifying association with the Major Facilitator Superfamily.

### Comparison of predictions made by Danchin and Fang

Recently Danchin and Fang^15^ used an “engineering-based” approach to investigate the unknown functions within the minimal bacterial genome and provided annotations for 58 and 13 proteins in the *generic* and *unknown* classes respectively. They set out to identify functions that would be expected to be in a minimal genome but were missing from the existing annotation and to then identify proteins that could perform these functions (although it is not clear how these candidates were identified as no methods were provided^15^).

Comparison of the results from both studies revealed considerable overlaps. Using our approach fourteen proteins remained in the *unknown* class without any assigned function, while Danchin and Fang did not provide any annotations for 26 and 52 proteins in the *generic* and *unknown* classes respectively. The predictions showed complete agreement for 34 proteins and minor differences for 18 proteins (Table S8). For a further 15 proteins the prediction was more detailed in one study than the other (Table S8). For example Danchin and Fang proposed that MMSYN1_0822, is an S component of ECF transporter and is part of a folate transporter, whereas we identified three possible folate transporters (MMSYN_0314, MMSYN_0822, MMSYN_0836) and could not confidently assign substrates to any of them.

Four of the predictions differed considerably (Table S8). They are represented by proteins such as MMSYN1_0388 which here was annotated as a transmembrane protein, possibly a cation transporter while Danchin and Fang suggested that it has a role in double strand break repair. For three of the proteins Danchin and Fang inferred more functional characteristics. They annotated MMSYN1_0853, MMSYN1_0530, MMSYN1_0511 with the functions energy-sensing regulator of translation, promiscuous phosphatase and double strand break repair protein respectively, while here they were retained as hypothetical since there was little agreement between the multiple methods used to be able to infer protein function.

## Discussion

The synthesis of a bacterium with a minimal bacterial genome resulted in an astounding number (149 of 473) of proteins of unknown function and emphasised the gaps in our understanding of the basic principles of life. We have demonstrated that many of these proteins of unknown function i) typically do not contain known protein domains ii) typically lack homology to proteins with known structure, and iii) are enriched for transmembrane proteins, many of which are likely to be transporters. Further, most proteins in the *unknown* confidence class appear to be bacteria- and probably clade-specific. These are highlighted by the 50 proteins for which the biological process is still classed as unclear and in particular for the 14 proteins that remain hypothetical with completely unknown function.

With the expanded functional assignments, 50% of the proteins encoded by the minimal genome perform functions associated with two fundamental life processes; *preserving and expressing genetic information* (Figure 3). Most notably many proteins were assigned transporter functions, and these proteins now represent 22% of the minimal genome. In generating the minimal genome, 32 *M.mycoides* genes with membrane transport functions, including several ABC transporters, PTS system proteins, amino acid permeases, Major Facilitator Superfamily proteins, were removed.^3^ Given that the experiment also removed many proteins with metabolic functions, the minimal genome bacterium is therefore reliant on obtaining many nutrients from the medium, and also the need to remove from the cell toxic molecules that may be generated. Thus it may not be surprising that transporters are essential for the bacterium. It was not possible to assign substrates for these transporters and it has been suggested that the mycoplasmal transport systems have broad specificity.

Our results demonstrate that the combination of a range of complementary advanced methods for protein function inference is superior to the use of individual approaches in the assignment of function for these ‘difficult’ proteins for which there is limited preexisting knowledge. Using a combination of results from 16 different methods we were able to reduce the number of proteins in the *unknown* confidence class to 41. Although, this approach was successful in general, there are still limitations and a need for further improvement. While functions have been assigned to many of the proteins that were previously unclassified, the assignments are often limited to general functions. Most changes of confidence elevated the classification from *unknown* to *generic*. For some proteins, more detailed functions were predicted by some of the methods, however in manually combining the predictions, there was insufficient evidence to assign them to a category higher than the *putative* class. Nevertheless, these functions should be sufficient to direct further research and experimental characterisation.

There is clearly a need for the development of additional approaches. Although multiple methods were combined to infer functions, many of these methods were homology based. However, most of the proteins of unknown function were homologous to few proteins with known functions and also lacked orthologs. Thus for many of the proteins where functions have been assigned, methods that are not dependent on homology were prevalent (e.g. FFPred^16^, Figure S4). This highlights the importance of developing further methods that do not rely on homology.

Comparison of our approach with that used by Danchin and Fang^15^ revealed considerable overlaps but also that our approach annotated many more proteins. It is difficult to expand on this comparison due to the lack of detailed methods provided by Danchin and Fang. It seems that their approach of looking for functions that appear to be currently missing from the annotated proteins in the minimal genome; requires significant knowledge of likely essential functions in a minimal cell, assumes that we know all of the essential functions within such a cell, and is further likely to exclude other functions that may not be considered essential in a minimal genome. In contrast, our approach sought to assign function to as many proteins as possible and then consider how these functions may relate to the minimal genome.

In summary, we successfully applied a combined bioinformatics approach to characterise the proteins with unknown function from the minimal genome that had not been annotated by previous approaches. Our approach utilised many state of the art methods including those that do not use homology. This has identified that a considerable proportion of the newly annotated proteins probably have transporter functions. These transporters are likely to be involved in the uptake of nutrients and efflux of waste products in a minimal genome organism that lacks many metabolic enzymes. Additionally, we identified that many of the unknown proteins were difficult to classify due to the limited information available about them (lack of orthologues and proteins domains and homology to known protein structures). Hence, we identified a need for additional, complementary approaches that enable assignment of functions to such proteins. Due to the physical limitations of the detailed experimental analysis of protein function on a large scale, such approaches will be critical for progress in synthetic biology and biotechnology applications.

## Methods

### Identifying basic protein properties

Protein domains were determined by running PfamScan against the library of Pfam 30.0 HMMs^10^. GO terms associated with Pfam domains were extracted using the pfam2go file^10^ (version 11 February 2017). The e-value of the domain matches were used to indicate the confidence of a GO term describing the function of the query protein. GO terms were clustered using REViGO^17^.

Orthologs were identified using EggNOG-mapper^18^ against HMM databases for the three kingdoms of life. Additionally, precision of predictions was prioritised by restricting results to only one-to-one orthologs. The EggNOG-mapper API was used to predict the orthologous groups in EggNOG that the minimal genome proteins belonged to. The proteins present in these orthologous groups were extracted and the species associated with the sequences were mapped to the NCBI Taxonomy to group them into phyla and used to identify the phyla where orthologues were present. Predicted features including GO terms, KEGG pathways and functional categories of Cluster of Orthologous Groups were also obtained from EggNOG-mapper.

### Identifying Membrane transporters and Lipoproteins

Proteins were classified as lipoproteins (SPaseI-cleaved proteins), SPaseI-cleaved proteins, cytoplasmic and transmembrane proteins using LipoP^19^. Similarly, proteins were distinguished between membrane transporters and non-transporters using TrSSP^20^ and SCMMTP^14^. TrSSP predicted substrates of the proteins from seven groups: amino acid, anion, cation, electron, protein/mRNA, sugar and other. The functions of membrane transporters and lipoproteins were further supported by identifying transmembrane helices, signal peptides and protein topology using TMHMM^21^.

### Inferring Gene Ontology based protein function

These methods included FFPred^16^, CombFunc^22,23^, Argot^24^ and LocTree3^25^ and CATH structural domains and functional families ^26^. Further functional information was extracted from the initial protein characterisation methods using TIGRFAM families of equivalogs, Pfam domains^10^, eggNOG orthologous groups ^18^, the top BLAST hit from UniProt and the protein structure modelled by Phyre2^11^. Additionally, where Phyre2 generated a confident model the structural methods firestar^27^, 3DLigandSite^28^and ProFunc^29^ were used. Given the large number of proteins in the generic and unknown groups that were predicted to be transmembrane, TrSSP^20^ and SCMMTP^14^ were used to infer transporter functions and lipoproteins were identified using LipoP ^19^.

GO terms were predicted using FFPred3^16^, Argot2.5^24^, CombFunc^22,30^ (only Molecular Function terms) and LocTree3^25^ (only Celullar Component terms). As the FFPred3 SVMs were trained only on human proteins from UniProtKB, predicted GO terms were additionally filtered using the frequency of terms in UniProtKB-GOA (version 5 June 2017). Predicted GO terms that were not annotated to any bacterial proteins in UniProtKB-GOA were removed from the set of FFPRED3 predicted functions as they were likely to be functions that are not present in prokaryotes.

Argot2.5 was run with the taxonomic constraints option. As scores returned by Argot2.5 have a minimum score of zero and no upper bound, the linear spline function recommended by the method developers (personal communication) was applied to rescale them to the range of 0 to 1. CombFunc ^22^ was run using standard settings.

### Structural Analysis

The CATH FunFHMMer web server was used to identify the functional families of structural domains, CATH FunFams.^26,31^ Where results did not identify a match to a FunFam, matches to CATH domains were considered. Concurrently, Gene3D^13^ was used to identify GO terms associated with CATH FunFams as well as to proteins with a similar multi domain architecture. Since UniProt accession codes were not provided for the minimal cell proteins, their closest UniProt homologs were determined by running BLAST^32^ against UniProtKB (version October 2016).^6^

Protein disorder was predicted using Disopred3.^33^ For each of the proteins, the percentage of disordered regions was calculated based on the Disopred3 results. Firestar^27^ and 3DLigandSite^28^ were used to predict ligands binding to the proteins. For Firestar only results marked as cognate were considered. Phyre2^11^ was run using standard mode to model the structure of the minimal genome proteins. Information provided by the name and description of the best matching models was used in the process of inferring function of the proteins. To make sure that each residue was covered with the highest possible confidence, the matches were firstly sorted by e-value and then selected gradually if they covered residues that were not covered before by a match with lower e-value. Structural models with at least 80% confidence were submitted to ProFunc^29^ to predict protein function from structure.

### Identifying operons

Genes in the synthetic *M. mycoides* (JCVI-syn1.0) were grouped into operons based on the predictions made for both *M. mycoides subsp capri LC* str 95010 and *M. mycoides* subsp mycoides SC str PG1 by two methods DOOR2^34^ and MicrobesOnline^35^. The proteins of the synthetic *M. mycoides* were first mapped to the proteins *of M. mycoides subsp capri* LC str 95010 and *M. mycoides* subsp mycoides SC str PG1 downloaded from GenBank. ^36^ This was done by using BLAST to search against databases constructed from proteomes of these two species and extracting the best hit. A protein from *M. mycoides* subsp capri LC str 95010 or *M. mycoides* subsp mycoides SC str PG1 was considered a corresponding homolog of a protein from synthetic *M. mycoides* if the coverage and identity were greater than or equal to 80%. Via the corresponding homologs, operons predicted for these two species by DOOR2 and MicrobesOnline were mapped to the proteins of the synthetic *M. mycoides*.

### Combined Protein Function Prediction

The results from the following methods were removed from the analysis if their e-value was above 0.001: TIRGFAM, Pfam, eggNOG-mapper, CathDB FunFams and domains. Models predicted by Phyre2 were kept if the probability of the match was above 80%, only results from Firestar marked as cognitive were retained. ProFunc results were not considered if scored as “long shots”. The best BLAST hit from UniProt was used to identify a UniProt identifier for the protein which was used for submission to Gene3D. Additionally, all the predictions of Gene Ontology terms were combined together and the probability of particular terms being predicted by any of the methods were calculated using the following formula:

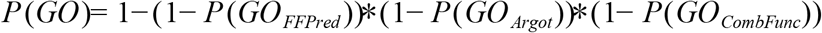

Only the best and consistently predicted Gene Ontology terms were examined for each of the proteins. For the final prediction of protein function results from all the methods were manually reviewed. The initial proposition of protein function was based on combining the results from TIGRFAM equivalog families, Pfam domains, eggNOG orthologous groups, CATH structural domains and functional families, the best BLAST hit from UniProt and Phyre2 model of structure. It was then verified using the best Gene Ontology terms, information on predicted ligands (Firestar, ProFunc, 3DLigandSite^28^) and transmembrane helices (TMHMM). Transporters and lipoproteins was predicted using membrane transporter (TrSSP, SCMTTP) and lipoprotein signal sequences (LipoP) respectively. Finally, it was considered if the predicted function was consistent within a group of genes in the same operon. If a specific function including a substrate and a biological role was determined and confirmed by multiple sources of evidence, a protein was classified in the *putative* confidence class. Similarly, if a general function was predicted and supported by several lines of evidence, a protein was classified in the generic functional class. Functional category was assigned to a protein if they were indicated coherently by any of the methods, e.g. if one method pointed confidently to transport and the other to proteolysis, the functional category remained Unclear.

## Authors’ contributions

MA and MNW devised the study. MA performed the experiments. All authors analysed the results and wrote the paper.

## Acknowledgments

The authors would like to thank Christopher Mulligan for helpful discussion about transporters, particularly ABC transporters and Prof. Stefano Toppo for advice regarding transformation of Argot2.5 scores into probabilities.

## Supplementary Figure Legends

**Figure S1.** Orthologs to the proteins in the minimal bacterial genome. The number of orthologs for each proteins identified in A) archea and B) eurkaryota. C) Summary of the total number of orthologs identified across different phyla for each of the functional confidence groups.

**Figure S2.** Confidence of the top structural template identified by Phyre2. The confidence score (0-100) is shown for the top scoring template identified for each of the proteins in the minimal genome. The score indicates the confidence that the template protein sequence and the minimal genome protein sequence are homologs.

**Figure S3.** Examples of proteins in the minimal bacterial genome that where it was difficult to predict their function. A)Protein MMSYN_0138 was previously completely uncharacterised and listed as a hypothetical proteins. Predictions for MMSYN_0138 by multiple methods identify a relationship to ATP binding domains of ABC transporters but the functional residues involved in ATP binding are not conserved making this function less likely. B) Protein MMSYN_0615 was previously classified as a tRNA binding protein in the *generic* confidence class. Multiple predictions suggest that it could be a Phenylalanine-tRNA ligase β subunit, however the β subunit in other bacteria typically contains around 800 residues, whereas MMSYN_0615 is only 202 residues. It therefore seems that tRNA binding is likely but the role of this function is not known.

**Figure S4.** Predictions made by the different individual methods used to infer functions of the minimal genome. For each protein in the *unknown* and *generic* confidence classes the methods that made predictions are shown. In each group, darker colours indicate that confident results were obtained from a method, whereas light colours indicate that a method did not provide a result that was useful. A) predictions from each method for proteins reclassified from the *generic* to *putative* class (on the left) and from the *unknown* to the *generic* class (on the right). B) the results from each method for proteins that remained in the *generic* (on the left) and the *unknown* classes.

## List of Supplementary Tables

**Table S1. Orthologs of the minimal genome proteins identified using eggnog**.

**Table S2. Domains identified in the minimal genome proteins.** Results are shown for search against the Pfam database of protein families.

**Table S3. Structural modelling of the minimal genome proteins.** Results are shown for modelling of the proteins using the Phyre2 server.

**Table S4. Membrane protein predictions for the minimal genome proteins.**

**Table S5. Inferred functions of the proteins encoded by the minimal bacterial genome.** The original annotation and the predicted functions from the analysis performed here are shown.

**Table S6. Gene Ontology based function predictions for the proteins encoded by the minimal genome.**

**Table S7. Gene3D predictions for the proteins encoded by the minimal bacterial genome.**

**Table S8. Comparison of the predicted functions of the minimal genome proteins with predictions made by Danchin and Fang.**

